# Extrinsic noise acts to lower protein production at higher translation initiation rates

**DOI:** 10.1101/2020.08.21.260976

**Authors:** Rati Sharma

## Abstract

Any cellular process at the microscopic level is governed by both extrinsic and intrinsic noise. In this article, we incorporate extrinsic noise in a model of mRNA translation and carry out stochastic simulations of the same. We then evaluate various statistics related to the residence time of the ribosome on the mRNA and subsequent protein production. We also study the effect of slow codons. From our simulations, we show that noise in the translation initiation rate rather than the translation termination rate acts to significantly broaden the distribution of mRNA residence times near the membrane. Further, the presence of slow codons acts to increase the mean residence times. However, this increase also depends on the number and position of the slow codons on the lattice. We also show that the the slow codons act to mask any effect from the extrinsic noise themselves. Our results have implications towards a better understanding of the role the individual components play during the translation process.

## I. INTRODUCTION

Despite having the same genetic information, the phenotypes of isogenic cells are not the same due to a variation in the protein expression among them. This variation is due to a combination of intrinsic and extrinsic fluctuations that influence the expression of proteins [1– 3]. In a biochemical system that consists of multiple reactants and products, the stochasticity in the system increases multiple folds. This results in an inherent randomness known as Intrinsic Noise (IN). The environment in which these biochemical reactions occur, introduces multiple stochastic variables (such as shared resources among populations of cells or genetic circuits and promoter architecture or pathways upstream of the circuit of interest, among others) that contribute to the Extrinsic Noise (EN) [2, 4, 5]. Noisy gene expression is known to be beneficial under conditions that are stressful or are prone to change. This lets a cell hedge its bet and adapt to changing scenarios [6–10]. Study of the effects of noise on gene expression can therefore lead to a better understanding of the phenotypic variability in these systems.

The fluctuations in protein expression can occur at different stages of the central dogma such as transcription, translation and post translational modification [1, 2, 11– 14]. In this work, we focus on the stochasticity inherent in mRNA translation, which itself is a complex process. The translation process broadly consists of three main stages, viz. initiation, elongation and termination. The ribosome machinery binds to the 5’ end of the mRNA during the translation inititation stage. This ribosome assembly then moves from codon to codon on the mRNA, in the process, adding amino acids to the growing oligomeric chain. This is the elongation stage. Finally, ribosomes exit the mRNA at the 3’ end, terminating the process of translation [15].

Even though the basic bits and pieces of central dogma have already been constructed, mRNA translation is still an active area of research. Several experimental studies are now looking at how the individual components of translation affect and aid protein synthesis [15–21]. These studies have also inspired several modeling frameworks that look at mRNA translation as a whole [22– 28], effects of the assembly of the ribosomal machinery [25, 29] or protein expression bursts as a result of stochastic transcription and translation [2, 4, 11, 30–40]. In this work, we focus on the kinetic aspects of mRNA translation. Specifically, like any cellular process, the three stages of translation are stochastic in nature and in the context of the translation process account for intrinsic noise. However, the rates of the three stages can themselves be stochastically fluctuating, which has its basis in extrinsic noise (EN) [36, 38, 41].

Here, we study the effects of EN on mRNA translation, which in turn has an effect on the ribosome residence time of the mRNA and the number of proteins produced. Further, we also look at the effect of slow codons on these properties. In section II, we describe the TASEP framework used to model mRNA translation. Section III gives a brief background and mathematical framework of extrinsic noise. Sections IV and V discuss the results and conclusions of this study.

## II. TRANSLATION MODEL AND THEORETICAL BACKGROUND

In bacteria, the process of translation takes place in the cytoplasm with the help of diffusing ribosomes. Further, several experiments exploring the sub-cellular spatial organization of bacterial cells, have also led to the observation that there are certain proteins that bind to the membrane and localize in specific pockets [42–45]. The mRNAs that code for these membrane binding proteins lie very close to the inner membrane of the bacterial cell. These observations lead to the hypothesis of cotranslational insertion [46, 47].

This system of cotranslational insertion has been theoretically studied using the Totally Asymmetric Exclusion Process (TASEP) framework [26]. This model is also one of the many that study ribosome dynamics on the mRNA, making use of the TASEP framework [22, 25, 27, 29]. The framework, in general, models unidirectional stochastic motion of particles on a one-dimensional lattice. The particles themselves are assumed to be point particles with the dynamical constraint that they can only hop on to the neighboring lattice site if it is unoccupied. Otherwise, the particles stay put where they are [48–51]. This system is therefore similar to the process of translation where ribosomes move from codon to codon on the mRNA from the 5’ end to the 3’ end in a unidirectional manner.

It is known that the translation initiation rate depends on the kinetics of the assembly of the ribosome machinery onto the mRNA, specifically on RNA unfolding kinetics and the affinity of a ribosome for the ribosome binding site [52]. In addition to this, the initiation rate is also prone to posttranscriptional regulation [53]. These can make the translation initiation rate dynamically fluctuating.

The translation elongation rates are also generally not uniform across codons on the mRNA [23, 54]. In fact, codons for which the tRNAs are rarely available in the cell tend to have ribosomes stuck on it for much longer than normal. These codons, termed slow codons, cause subsequent slowing down of the protein synthesis not just for that particular ribosome but also for those that come after it [54]. These ribosomal traffic jams therefore lead to longer ribosomal residence times and changes in the gap distribution. As defined in an earlier study [26], we consider the average ratio of the rates *α* to *γ* to vary between 0.01 to 0.4. Also, we set the translation elongation rate (*γ*) to be equal to the the translation termination rate (*β*) as they are generally of the same order [52, 53, 55].

In our study of extrinsic noise on mRNA translation, we use the TASEP framework [26] to model ribosome dynamics on the mRNA lattice. In this model, a ribosome binds to the mRNA with rate *α* at the 5’ end. The ribosome then moves on to the subsequent codons on the mRNA lattice with rate *γ*. Once the ribosome reaches the 3’ end, it exits the mRNA with rate *β*. Our model of the mRNA is assumed to have 100 lattice sites, each site accounting for 10 codons [26] (corresponding to 30 nucleotides, which is the average ribosomal footprint of *E. coli* [21]). TASEP models which include full extensions for ribosomal footprint and therefore incorporate exclusion interaction lengths between two ribosomes do not give significantly different results from the simpler model [56]. Therefore, similar to a recent study [26], we proceed to use the simple TASEP framework for our system.

We study the effects of EN on such a translation mechanism by incorporating fluctuating rates of translation initiation and termination. A schematic of this framework is depicted in Fig. 1. We carry out a Monte Carlo simulation study of this system to account for the effects of EN on ribosome translation and protein production. We also look at these effects in the presence of a single or multiple slow codons in the mRNA.

**Figure 1.**
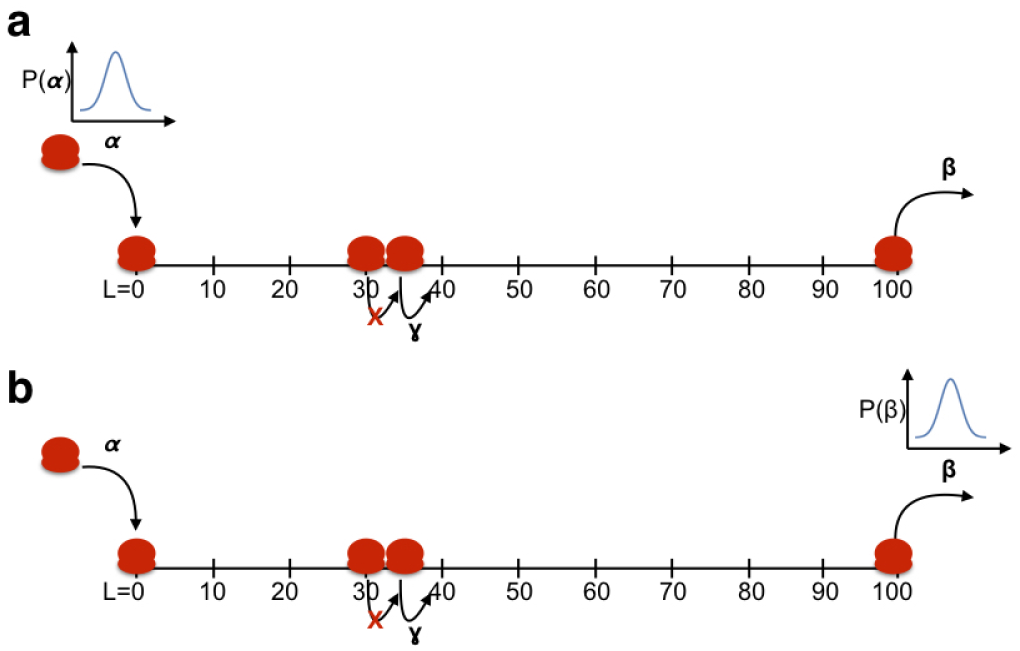
Extrinsic noise in the TASEP model of mRNA translation. Depiction of noise in the rates of (a) translation initiation (rate *α*) and (b) translation termination (rate *β*). The red blobs represent the ribosome assembly. A ribosome enters the mRNA from the 5’ end. It then moves from codon to codon (if the site is unoccupied) with rate *γ* in a unidirectional manner towards the 3’ end, in the process adding amino acids to the growing oligomeric chain. It exits the mRNA once it reaches the 3’ end.

In the next section, we describe the details of incorporating EN into the system.

## III. INCORPORATING EXTRINSIC NOISE INTO MRNA TRANSLATION

Extrinsic noise can be incorporated into the translation initiation rate (*α*) or the translation termination rate (*β*), by multiplying the kinetic rate constants with a random number picked from a previously specified distribution. In this work, we study the effect of EN through Ornstein-Uhlenbeck (OU) process, a common noise distribution used in several studies [33, 36, 57–59].

Ornstein-Uhlenbeck (OU) process is a stochastic Markov process (represented here by the function *ξ*(*t*)), whose dynamics is specified by the following differential equation.

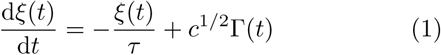

Here, *τ* is the relaxation time, Γ(*t*) is the Gaussian white noise and *c* is the diffusion constant given by *c* = 2*D/τ*.

D is a measure of the noise strength, which specifies the deviation of the noise from its mean. The OU process has an exponential time autocorrelation function given by

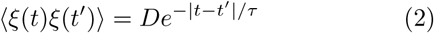

Thus, *ξ*(*t*) and *ξ*(*t*^′^) lose correlation when |*t*−*t*^′^| is greater than *τ*. Dynamics of *ξ*(*t*) is simulated using the following discrete equation [33].

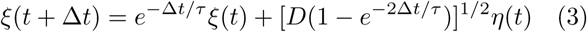

where *η*(*t*) is a unit normal random variable. In our simulations, we start with the initial condition of *ξ*(0) = 0, which leads to ⟨*ξ*⟩ = 0, where the angular brackets denote an average over all realizations of the noise. We also carry out all our simulations with Δ*t* = 0.01*s*, which is much smaller than or equal to the the value of *τ* used to incorporate EN.

In our study, at each time step of the simulation, *ξ*(*t*) is picked from the OU process for specific values of *τ* and *D*. Subsequently, EN is incorporated into rate constants by replacing *α* and *β* with *αe*^*ξ*(*t*)^ and *βe*^*ξ*(*t*)^ respectively. The intensity of the noise is varied by varying *τ* and *D*. A higher *τ* corresponds to higher correlation time leading to an increase in EN. Similarly, a high *D* corresponds to a broader distribution (increased variance) of EN, thereby increasing the chances of higher rates being incorporated into the system.

We carry out Monte Carlo simulations of our TASEP model without extrinsic noise using the well known Gillespie algorithm [60]. The simulations of the model in the presence of extrinsic noise are carried out using the EXTRANDE algorithm, which is a modified version of the Gillespie algorithm, developed to simulate a network in a dynamically fluctuating environment [61].

## IV. RESULTS

### A. Ribosome residence times

As a first step, we study the effect of EN on the residence time distribution of the ribosome along the mRNA lattice. It has been shown that the residence time on a lattice of length *l* follows a gamma distribution [26], given by,

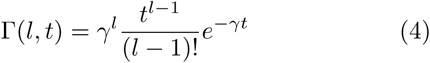

where *γ* is the time correlation constant with dimensions *s*^−1^. In Figs. 2(a and c), we show the EN distributions for two different values of *D*, while keeping *τ* fixed. These EN distributions are then incorporated into the kinetic rates of initiation or termination to obtain the ribosome residence time distributions from simulations of the ribosome trafficking. These are shown in Figs. 2(b and d). From the simulations, it is observed that noise in *α* broadens the residence time distribution compared to the no EN condition, whereas, the distribution for the case when the noise is incorporated in *β* is similar to the no EN condition. One can also observe deviations from the gamma distribution when EN is incorporated in *α*. Since EN in *α* shows greater effect on residence times of ribosomes, most of the following analyses are carried out with stochastically fluctuating *α*.

**Figure 2.**
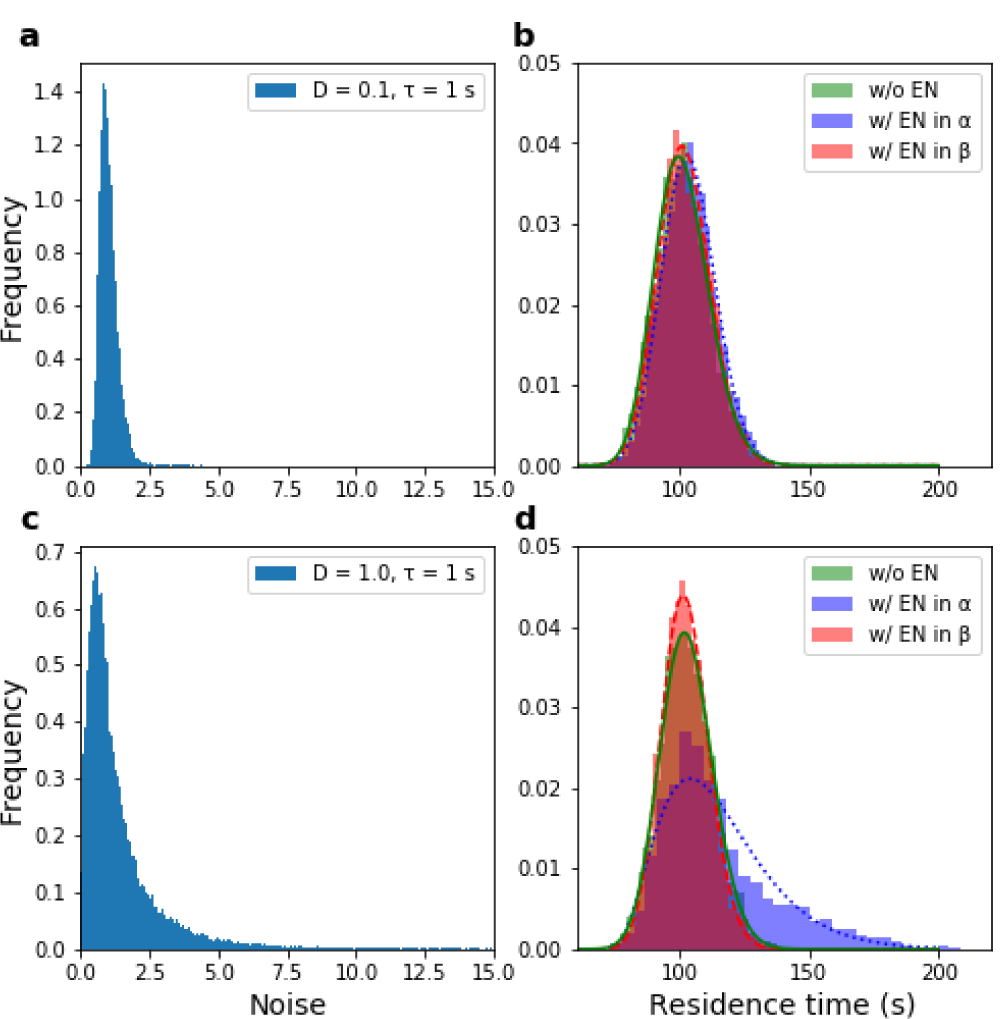
Effect of EN on ribosome residence times. (a) Simulated OU noise distribution that is incorporated either in the translation initiation rate (*α*) or the translation termination rate (*β*) with *D* = 0.1. (b) Distribution of residence times of ribosomes on the mRNA lattice without EN (green) and when incorporating EN from an OU process in the translation initiation rate, *α* (blue) and the translation termination rate, *β* (red) for the same values of *D* as in (a). (c and d) Same as (a and b) but for *D* = 0.1. All the other parameters are fixed at *τ* = 1.0*s, < α >*= 0.4*s*^−1^, *< γ >*= 1.0*s*^−1^ and *< β >*= 1.0*s*^−1^. Histograms are obtained from simulations and solid lines are fits to gamma distributions. The overlaps in the distributions are shaded darker and in a different colour.

### B. Exploring specific effects of the EN parameters

Protein production from mRNA depends on the amount of time a ribosome takes to traverse the complete mRNA and finally exit it at the 3’ end. Through our simulations, we calculate this ribosome residence time, distributions of which are shown in Fig. 2. Production of a single complete protein is accounted for upon the exit of a ribosome from the mRNA.

In order to discern the effect of the two parameters of EN (*τ* and *D*) on the translation process, we compare the ribosome residence times and the protein expression trends with those obtained for the no EN case. These are shown in Figs. 3 and 4. The simulations for these were carried out up to a maximum time of 8 minutes (which is the average lifetime of mRNAs in E. coli [62]) in each case. This allows for direct comparison of the number of proteins produced at the end of the simulation time. The curves show the mean and the standard error over mean (SEM) calculated from 100 such simulation runs.

**Figure 3.**
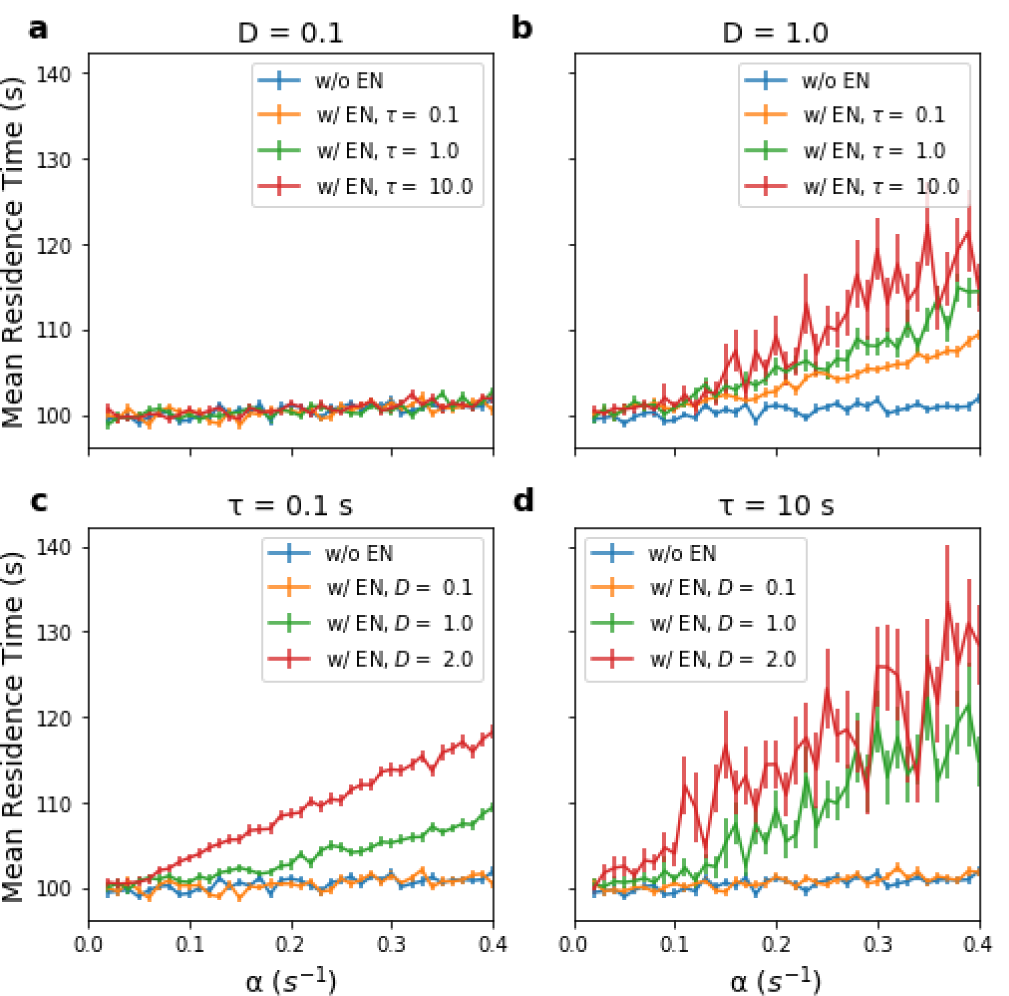
Ribosome residence times vs. *α*. Plots of mean ribosome residence times on the entire mRNA lattice versus increasing translation initiation rates (*α*) [Top row:] Plots are for (a) D = 0.1 and (b) D = 1.0 at fixed values of correlation times, *τ* = 0.1*s* (orange), *τ* = 1.0*s* (green), *τ* = 10*s* (red). [Bottom row:] Same as top row for (c) *τ* = 0.1*s* and (d) *τ* = 10.0*s* for fixed values of diffusive strengths, D = 0.1 (orange), D = 1.0 (green) and D = 2.0 (red). All the plots are compared to that obtained with no EN in the translation initiation rate, *α* (blue). All the other parameters are fixed at *β* = 1.0*s*^−1^ and *γ* = 1.0*s*^−1^.

**Figure 4.**
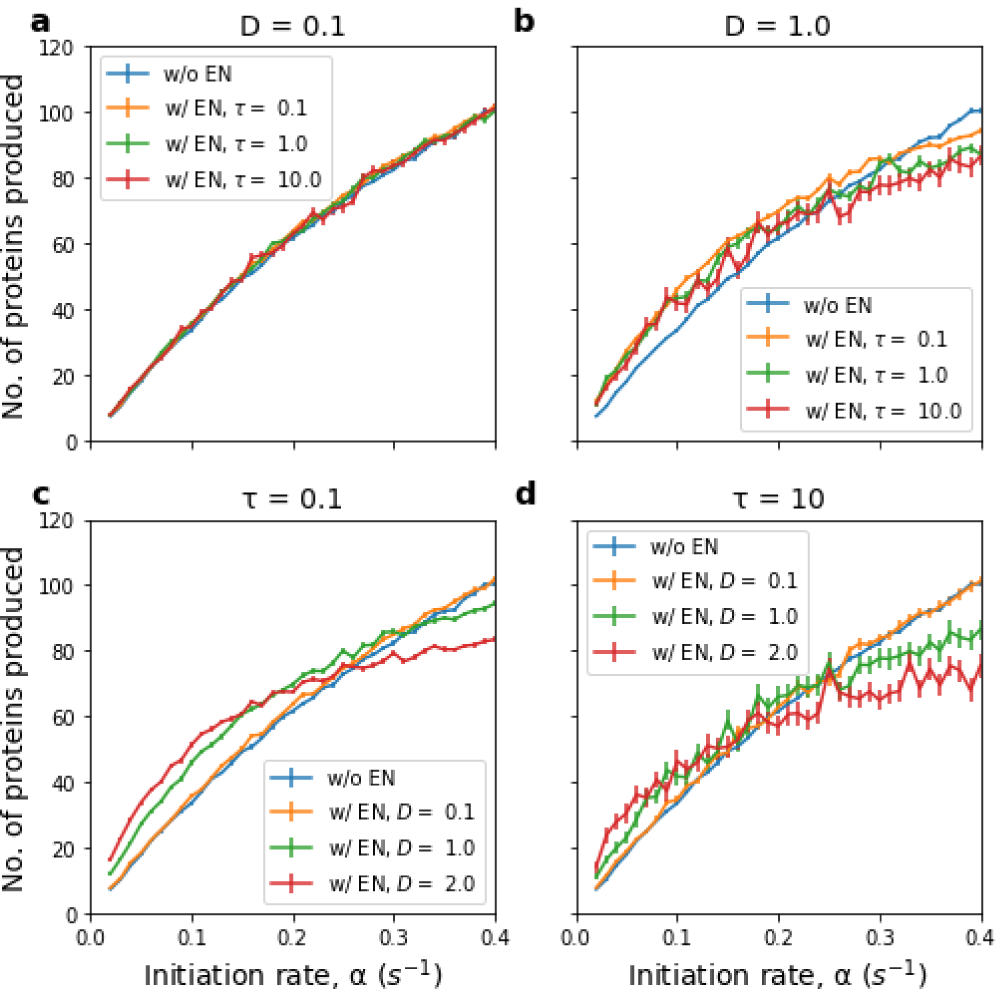
Protein expression vs. *α*. Plots of mean number of proteins produced versus increasing translation initiation rate (*α*) [Top row:] Plots are for (a) D = 0.1 and (b) D = 1.0 at fixed values of correlation times, *τ* = 0.1*s* (orange), *τ* = 1.0*s* (green), *τ* = 10*s* (red). [Bottom row:] Same as top row for (c) *τ* = 0.1*s* and (d) *τ* = 10.0*s* for fixed values of diffusive strengths, D = 0.1 (orange), D = 1.0 (green) and D = 2.0 (red). All the plots are compared to that obtained with no EN in the translation initiation rate, *α* (blue). All the other parameters are fixed at *β* = 1.0*s*^−1^ and *γ* = 1.0*s*^−1^.

These plots clearly show that for low diffusive strength (*D* = 0.1, Fig. 3a), the mean residence time increases slowly (non-negative slope greater than 1) with increasing *α*. Further, with this low *D*, the curves are equivalent to those obtained in the no EN case even for higher *τ* values. The effect of EN manifests itself in the case of high *D* (*D* = 1.0, Fig.3b), even when *τ* is low (*τ* = 0.1*s*). This is also reflected in Fig. 3c. However, as evident from Fig. 3(d), high *τ* combined with high *D* leads to increased variation in the residence time.

We next compute the total number of proteins produced at the end of 8 minutes of real time in the simulation. For low *D* (Fig. 4a), the number of proteins produced vs. *α* follows the same trend as the no EN case. However, for high *D* (Figs. 4b, c and d), protein production increases with increasing *α* up to a point and then starts to flatten. This is a result of two opposing forces – (i) increase in translation rate due to increasing *α*, which tends to raise protein production and (ii) an increase in the ribosome residence time (a consequence of increasing EN), which tends to decrease protein production. As evident from Figs. 4(b, c and d), the slope change (flattening) in these scenarios happens at successively lower *α* with increasing EN.

### C. Ribosome traffic in the presence of slow codons

As mentioned earlier, many mRNAs invariably have certain codons or sequences of codons that slow down ribosome trafficking at that site. The effects of these slow codons are long lasting. We now explore their effects on mRNA translation, protein production and ribosome trafficking.

We first look at the effect of the lattice site position of the slow codon on the mRNA. Referring to Fig. 1, the mRNA is considered to be composed of 100 lattice sites (l), with 1 representing the first site from the start codon and 100 the farthest on the right. With this picture, one can now see in Fig. 5, that the ribosome residence time increases linearly with the slow codon site. This can be a manifestation of the fact that the farther the slow codon site, the greater the traffic jam experienced by the ribosome on the mRNA lattice. Further, it seems that EN has no effect on the linear increase shown here when the elongation rate, *γ*, is decreased to 1 tenth the usual rate.

**Figure 5.**
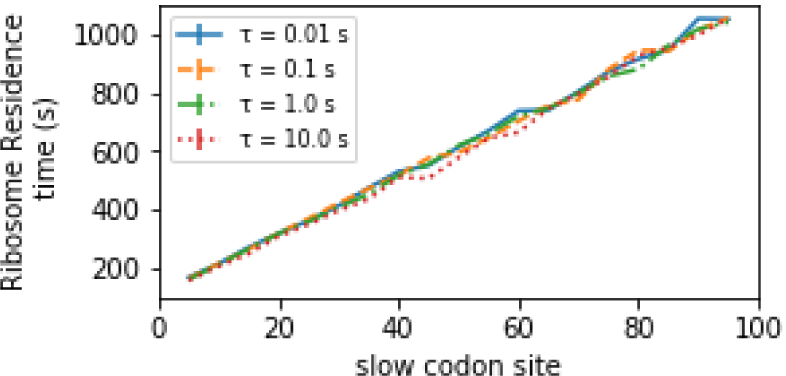
Effect of slow codons. Ribosome residence time versus the slow codon site (l) on the mRNA lattice for *γ* = 0.1*s*^−1^ at the slow codon site, *γ* = 1.0*s*^−1^ at all other sites, *β* = 1.0*s*^−1^, *D* = 1.0, and *τ* = 0.01*s* (blue solid line), 0.1*s* (orange dashed line), 1.0*s* (green dot dashed line) and 10*s* (red dotted line).

In the simulations represented by Fig. 5, we only considered one slow codon. We now also consider two slow codons placed 20 lattice sites apart and look at the ribosome trajectory and gaps. This is shown in Fig. 6. The slow codon sites are evident in the ribosome trajectories by the build up of ribosomes below that site. One can also observe that the build-up gets denser with two slow codons compared to just the one. This build-up is also reflected in the gap distribution among the three cases. mRNAs with no slow codons have a gap distribution that tapers off only at 15-20 lattice sites, whereas the positions before a slow codon have a gap no more than a single lattice site. The distribution of gaps broadens again beyond the slow codon, however it is more long-tailed than the no slow codon case. Finally, the trajectories beyond the two slow codons show lesser broadening in the gap distribution compared to the no slow codon case. Surprisingly, the no EN and with EN scenarios for the slow codon cases do not show any major difference between them. This could be because the slow codon masks the effect of EN.

**Figure 6.**
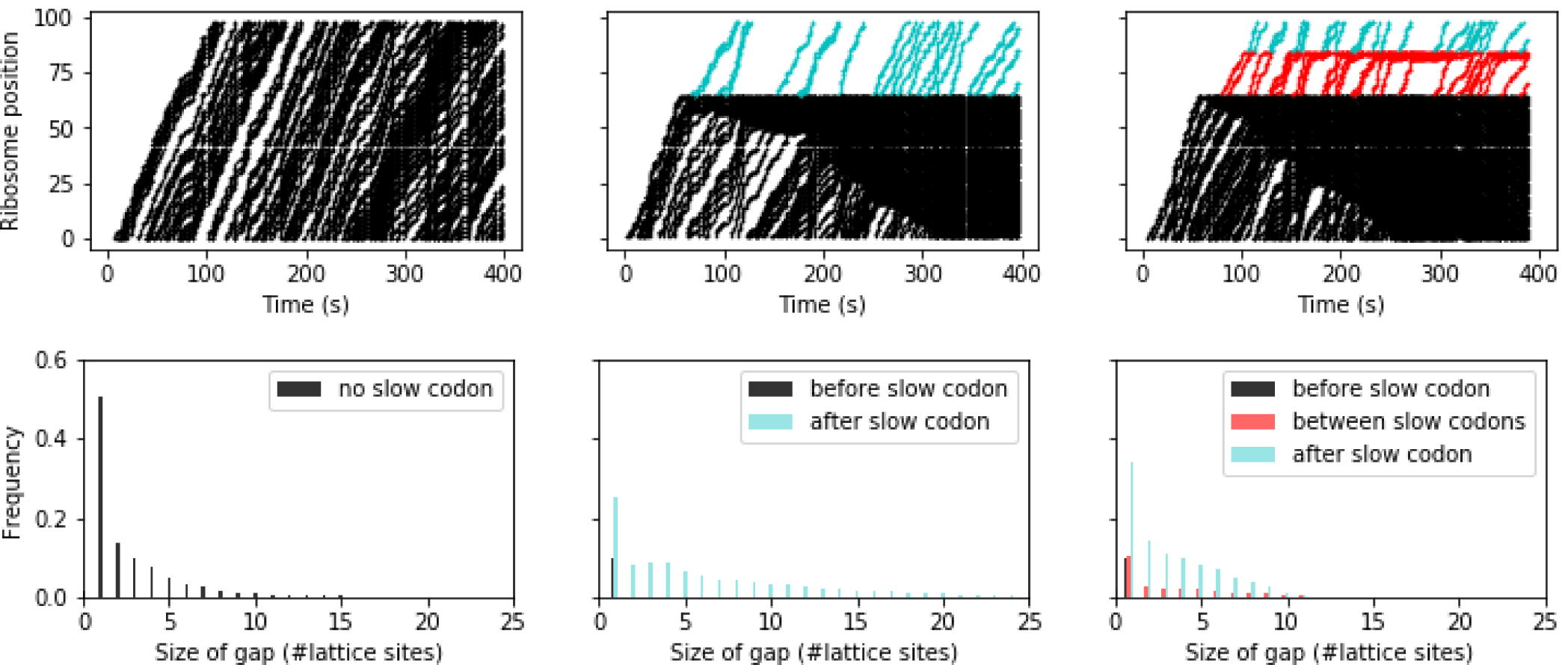
Ribosome trafficking. [Top:] Ribosomal position versus time and [Bottom:] frequency versus gap length between successive ribosomes on an mRNA with (a) no slow codons, (b) 1 slow codon at site *l* = 65 and (c) two slow codons with the second slow codon site at *l* = 85. Ribosomes and gaps before the first codon are shown in black, after the slow codons are shown in cyan and between two slow codons are shown in red. All the other parameters are fixed at *α* = 0.4*s*^−1^ *β* = 1.0*s*^−1^, *γ* = 0.1*s*^−1^ for the slow codon site, *γ* = 1.0*s*^−1^ for all other sites, *D* = 1.0 and *τ* = 1.0*s*.

## V. CONCLUSIONS

### EN in *β*

Since translation termination (kinetic rate *β*) is a rapid event compared to translation initiation (kinetic rate *α*), as shown in Fig. 3, the order of magnitude of the noise that affects the rate *α* does nothing to perturb *β*. Of course, if the rate *β* is made comparable to that of rate *α* = 0.4*s*^−1^, the ribosome residence time distributions show deviation from the gamma distribution. Not only that, in this scenario, the distribution obtained for the case of EN in beta is shifted to the left compared to the no EN case. Also, all the distributions are long tailed. This is shown in Figs. 7(a and b). However, when *β* = 10.0*s*^−1^ *>> α*, as in Figs. 7(c and d), there is virtually no difference between the ribosome residence time distributions for the three cases, viz. no EN, EN in *α* and EN in *β*. The distributions in Figs. 7(c and d) also follow the gamma distributions. This analysis shows that the ratio between the three rates plays a major role in determining the nature of the residence time distributions. This then is also liable to affect protein production and gap distribution.

**Figure 7.**
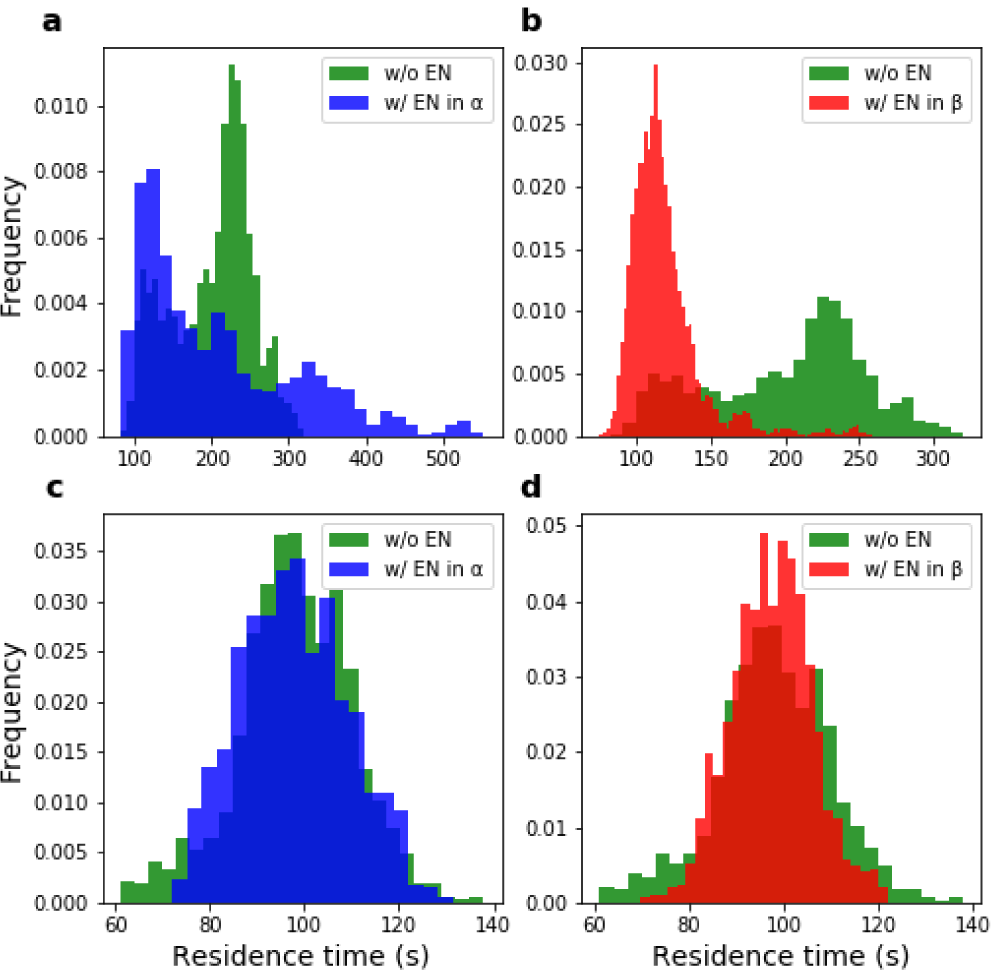
Effect of changing alpha to beta ratio on ribosome residence times. [Top:] Comparison of ribosome residence time distribution obtained from simulations with no EN and when EN is incorporated either in (a) the translation initiation rate (*α*) or (b)the translation termination rate (*β*) with *< β >*= 0.4*s*^−1^. [Bottom:] Same as top with *< β >*= 10.0*s*^−1^. All the other parameters are fixed at *D* = 1.0, *τ* = 1.0*s, < α >*= 0.4*s*^−1^ and *< γ >*= 1.0*s*^−1^. Histograms are obtained from simulations. The overlaps in the distributions are shaded darker.

### Protein production

In elucidating the effects of EN on mRNA translation, we see that the protein production shows a gradual increase with *α* (Fig. 4). This gradual increase also has two regimes, categorized by the slopes (high and low). For the low EN case (Fig. 4a), there is minimal change in the slope. However, for high EN (Fig. 4b, c and d), the change in slope is more pronounced. Further, a smooth transition is observed between the low and high slope regions of the curve. The property of gradual slope change is similar to the slope change observed in the ribosomal current in a previous study on mRNA translation in the yeast strain, *S. cerevisiae* [23]. Since protein production is a direct consequence of ribosomal current, the trend shown here is in agreement with expected behavior. However, the change in slope becoming more pronounced for high EN is a useful property that came out of this study. Since, number of proteins produced and its distribution is a quantity that can be determined from experiments, these slope changes can also be used to determine the extent of the extrinsic noise in the translation initiation rate.

### Presence of slow codons

In our simulations, the slow codons act to mask the effect of EN, i.e., they raise the mean ribosome residence time to such an extent that there is virtually no difference between the simulations with and without EN. Also, as indicated in Ciandrini et al’s yeast analysis [23], the position of the slow codons does have an effect on the ribosomal current, which is attributed to a buildup of ribosomes before the slow codon site. This buildup manifests in the residence time distribution in our analysis. Here, we see in Fig. 5, that the buildup leads to a greater increase in the ribosome residence time even though there is no change in the rate. This buildup can also be seen in Fig. 6, which further changes the gap distribution profile.

Finally, this analysis of extrinsic noise in mRNA translation helps us understand and deduce the source of fluctuations in protein production. Specifically, under realistic conditions, noise in translation initiation has a major effect and through the analysis of protein production, one can even deduce the extent of that noise. Further, this analysis shows us that slow codons are also a source of fluctuation and the different gap distributions can be used to determine the presence of slow codons. One can also extend this study to include specific effects from diffusing ribosomes and drop off of these ribosomes from the mRNA lattice. These will be taken up in future studies. The analyses presented in this simulation study can also help in carrying out a similar analysis in experimental studies. This can then help in factoring in the effect of individual processes of mRNA translation that contribute to noisy protein distributions.

